# Evidence that genome editing is preferable to transgenesis for enhancing animal traits

**DOI:** 10.1101/2025.04.25.650727

**Authors:** Jinhai Wang, Shinichi Nakagawa, Jiaqi Wang, Robert Stewart, Alexandra Florea, Rex A. Dunham, Fei Ling, Gaoxue Wang, Lily Liu, Diego Robledo

## Abstract

Production traits such as growth, disease resistance, and fatty acid content in engineered animals are anticipated to be enhanced *via* transgenesis or genome editing. It is, however, unclear whether this expectation is upheld across taxa on a global comparison. Here, we perform a meta-analysis of 154 studies involving 72 species and 55 genes to quantify and compare the effects of transgenesis and gene editing on production traits in animals. While transgenesis is more commonly applied for trait enhancement, gene editing demonstrates more pronounced and widespread effects, particularly on growth and disease resistance traits, as reflected by larger effect sizes and broader impacts across trait responses. Yet, we observe taxon- and parameter-specific differences in patterns of trait enhancement. For instance, transgenesis reduces pathogen load in chickens and cattle but not in pigs, whereas gene editing lowers virus RNA levels in pigs but has limited success in chickens and cattle. In contrast, both transgenesis and gene editing significantly increase growth rates in ray-finned fishes. Notably, although transgenes or edited genes remain highly expressed or repressed in F_1_ offspring, the magnitude of trait improvement is diminished compared to the founder generations. This study provides evidence-based insights that can assist researchers in refining their methods and directing future investigations into trait enhancement in genetically engineered animals, while also playing a key role in policymaking.

## 1. Introduction

Genetic engineering (transgenesis) and gene editing have become cornerstone technologies in modern biology, providing tools to make precise alterations in the genomes of organisms. Techniques such as CRISPR/Cas9 [1], TALENs (Transcription Activator-Like Effector Nucleases) [2], and ZFNs (Zinc Finger Nucleases) [3] have been widely adopted for their ability to target specific genes associated with desirable traits in animals. These technologies have been applied to a wide range of species, from laboratory animals to livestock, poultry and fish, with the goal of enhancing desirable traits in animals, such as increased growth rates [4,5], improved disease resistance [6-8], and optimized fatty acid composition [9-11]. Moreover, these technologies are instrumental in biomedical research for generating animal models of human diseases [12,13], as well as in conservation biology to support the protection of endangered species [14,15].

Since the 1980s, transgenic farm animals have been investigated for their potential as bioreactors [16,17], capable of producing recombinant proteins. This investigation led to the introduction of a diversity of growth hormone (GH) and growth factor genes *via* transgenesis to enhance growth in livestock species, such as pigs [18,19] and sheep [20,21], as well as in fish, including Atlantic salmon (*Salmo salar*) [22,23], common carp (*Cyprinus carpio*) [24], Nile tilapia (*Oreochromis niloticus*) [25], and channel catfish (*Ictalurus punctatus*) [26]. Additionally, transgenic farm animals with enhanced disease tolerance and altered fatty acid profiles have been developed, encompassing both livestock and aquatic species [27,28]. Recent advancements in genome editing technologies, particularly CRISPR-led tools, have increasingly facilitated genetic modifications in farm animals. Nevertheless, to date, only *GH*-transgenic AquAdvantage salmon [29,30] and transgenic GalSafe pigs [31] have been approved for human consumption, with stringent regulatory oversight making it challenging for genetically modified organisms to enter the food supply. In contrast, gene editing can result in changes that mimic natural mutations, may face fewer regulatory obstacles than transgenesis in some countries [32,33]. Several gene-edited animals, including *mstn*-edited red sea bream and leptin receptor-deficient tiger puffer [34] have been approved by governments. There remains a critical need to communicate to policymakers the potential benefits of gene editing and genetic engineering in animal applications, given the advancements over recent decades. Although both methods can effectively improve desired traits, no comprehensive studies have systematically compared the relative efficacy of transgenesis and gene editing in improving these traits, apart from their distinct regulatory challenges.

Current evidence for up-scaling favorable traits *via* transgenesis or gene editing remains inconclusive, with independent studies yielding varied and, in some cases, conflicting conclusions. The effect of transgenesis or gene editing on trait improvement may vary by different parameters and animal taxa. For instance, Pursel et al. [19] reported that *IGF-I* (insulin-like growth factor I) transgenic pigs did not show improvement in body weight or specific growth rate compared to non-transgenic individuals. However, a subsequent study by Bee et al. [35] suggested that although *IGF-I* transgenic pigs had lower carcass dressing percentage, they did exhibit a larger longissimus dorsi muscle compared to sibling controls. Several studies showed that disruption of the myostatin (*mstn*) gene has been shown to improve growth by increasing myofiber numbers in pigs [36,37], sheep [5] and fishes [38], but not changing myofiber size. Conversely, other studies have reported increases in both myofiber size and number in *mstn*-mutant pigs [39], chickens [40] and fishes [41-43]. Additionally, transgenic channel catfish expressing lysozyme exhibited enhanced lysozyme activity following bacterial infection compared to non-transgenic controls [44], while no significant differences in serum lysozyme levels were observed between lactoferrin-transgenic and control grass carp (*Ctenopharyngodon idellus*) after bacterial challenge [45]. Documented investigations have shown that *CD163*-edited pigs had enhanced resistance against porcine reproductive and respiratory syndrome virus (PRRSV) [46-49], but not to the African swine fever virus [50]. Fatty acid-related traits have similarly demonstrated inconsistent outcomes. Transgenic pigs and cattle expressing the omega-3 desaturase (*fat1*) transgene did not show significant alteration in eicosapentaenoic acid (EPA), docosahexaenoic acid (DHA), or n-6 polyunsaturated fatty acid (ω-6) content [51,52], whereas other studies observed significant increases in EPA and DHA and reductions in ω-6 levels in *fat1*-transgenic pigs [53,54]. Additionally, studies on *elovl2*-transgenic channel catfish (*Ictalurus punctatus*) have reported both significant and non-significant changes in DHA, ω-3, and ω-6 fatty acid content, depending on the specific study [11,55].

The variability in the reported effects of transgenesis and genome editing on trait enhancement may arise from diverse biological and methodological factors that influence the outcomes of these interventions. A meta-analytical approach is essential to understand the influence of these ‘moderators’ and to explore the general effects on trait performance. As a result, the observed heterogeneity in trait performance enhancements reflects the specificity of these factors, which must be accounted for when evaluating the effectiveness of transgenesis or gene editing in improving traits. Yet, no study has done this systematically for animals. Addressing this gap requires a comprehensive integration of all available data and a re-quantitative analysis.

In this study, we aimed to investigate how potential determinants affect the improvement of trait performance using multiple moderator analyses based on a meta-data matrix. The primary objective was to quantitatively integrate empirical data on the application of transgenesis and genome editing for trait enhancement in farm animals. Specifically, we aim to (1) identify consistencies across studies involving transgenesis or gene editing in different animal taxa to evaluate the performance of transgenic and gene-edited animals in terms of gene/protein expression, growth, disease resistance and fatty acid profiles based on various parameters; (2) compare the effects of transgenics and gene editing on trait enhancement by different parameters to determine whether genome editing offers superior results compared to traditional gene transfer; (3) assess whether the impact of transgenesis and gene editing on trait improvement varies with the age of the engineered animals; and (4) determine promising candidate transgenes or innate genes capable of consistently improving traits across taxonomic classes. Herein, through a cross-taxon meta-analysis and a global synthesis of published data, we provide a comprehensive understanding of how to optimize transgene or innate gene manipulation to enhance trait performance in farm animals using genetic engineering or genome editing approaches.

## 2. Materials and methods

### 2.1. Literature search

A literature search of the *PubMed, Web of Science* and *SCOPUS* databases using search strings (*Note 1 in Appendix2*) yielded over 2,000 potentially eligible published articles from each database. The search and collection of literature were carried out based on the statement of Preferred Reporting Items for Systematic reviews and Meta-Analyses for Ecology and Evolutionary Biology (PRISMA-EcoEvo) [72]. We also conducted a forward search using 10 relevant review papers related to the topic of our meta-analysis [7,27,60,67,73-78]. The last retrieval date for these online databases was 17 September 2024. Initially, 10,299 peer-reviewed scientific articles were retrieved using the above strings (*Fig. S1 in Appendix2*). After a thorough review of the abstracts and full texts, we deemed 154 publications comprising a total of 1,145 data entries to build our meta-dataset (*Appendix1*). We did not request the unpublished datasets from colleagues due to the risk of biasing the estimates of effect size [79]. We ensured that the screening process was highly reproducible (*Note 2 in Appendix2*).

### 2.2. Selection criteria

To ensure the relevance and rigor of the meta-analysis, the following selection criteria were included: (1) the article should be a study-based research (not a meta-analysis, a review, or case study) on non-human animals, written in English; (2) peer-reviewed studies involving controlled experiments with genetically engineered or edited animals were included, while studies that were observational or theoretical were excluded; (3) recruited articles must have investigated quantitative data on at least one trait performance metric *in vivo*, such as growth, disease resistance, or fatty acid content in wild-type (WT, non-transgenic or non-edited) and treated (transgenic or gene-edited) groups to calculate effect sizes.

### 2.3. Information extraction and moderators

To determine gene/protein expression and quantify the evidence for the improvement of trait performance (growth, disease resistance, or fatty acid) through transgenesis or gene editing in animals, specific data were extracted from selected studies. This process included the following details: author information (first author, year), article title, taxon, species/breed from each taxon, method used (transgenesis or gene editing), target gene, and trait of interest, as well as sample size, means and standard deviation (SD) or standard error (SE) of outcome data in WT and treated groups (*Appendix1*). Where figures were the only source of data, we employed ImageJ.JS (https://ij.imjoy.io/) to derive data (mean and SD or SE).

To investigate the influence of methodological moderators on trait performance (*Note 3 in Appendix2*), we initially categorized the dataset into two groups based on the applied method (transgenesis and gene editing) using the method-moderator analysis. Subsequent moderator analyses were then conducted separately within each group. To understand how biological moderators affect trait enhancement, data on various biological variables were compiled from the 154 studies included in our meta-analysis. We began by evaluating the overall effects of transgenesis and gene editing on target gene or protein expression, capturing information on the taxon, species or breed, target gene, trait, and metrics for gene or protein expression for each study. Transgenesis, as is well-established, typically involves integrating transgenes into the genome, often under the control of specific promoters to drive overexpression [7]. Furthermore, gene or protein expression often displays tissue-specific patterns, as described in individual studies [78]. Consequently, we conducted moderator analyses stratified by taxon, species, gene, trait, promoter, and tissue to identify significant variations in gene or protein expression. Parallel analyses were conducted for the gene editing group, excluding promoter-moderator analyses, to evaluate the effects of these moderators on gene/protein expression.

With respect to growth traits, various measured parameters, including body weight (e.g. mean weight, average weight, carcass weight, wet weight, mean mass, live weight, dressing percentage), condition factor (e.g. condition score), feed conversion efficiency (FCE), myofiber number (e.g. fiber cell number, myofiber nuclei number, total number of fibers, percentage of fibers, myofiber density), and myofiber size (e.g. mean muscle fiber cross-sectional area, muscle fiber area, area of fiber cells, mean/total area of fibers). Additional measurements included plasma growth hormone (GH) (e.g. concentration of GH), specific growth rate (SGR) (e.g. average growth speed, standard growth rate, percent of body weight increase), weight gain (e.g. percentage weight gain, daily weight gain). The disease resistance traits were either measures of antibody response (e.g. ELIAS S/P ratio), cumulative survival rate (CSR), lysozyme activity, pathogen load (e.g. number of virus-positive cells, colony-forming unit, viral growth, number of surviving bacteria, viral particle production), phagocytic activity, sign score (e.g. score for clinical signs, score for pathological changes), virus RNA (e.g. relative expression of virus RNA, viral nucleic acid, fold change of viral RNA, viral RNA load level, viral RNA copies), or virus titer (Log10TCID50, mean viral shedding titer, number of plaques). Similarly, fatty acid traits were captured by measuring multiple parameters, including the content of docosahexaenoic acid (DHA), eicosapentaenoic acid (EPA), *n*-3 polyunsaturated fatty acid (ω-3), *n*-6 polyunsaturated fatty acid (ω-6), and the ratio of ω-6 to ω-3 (ω-6/ω-3) (*Note 4 in Appendix2*).

### 2.4. Effect-size calculations

The goal was to determine trait enhancement in transgenic and gene-edited animals relative to WT individuals, and to compare the effects of transgenesis and gene editing on trait enhancement across various parameters in animals. Herein, all outcome parameters were continuous variables, and sample sizes were not identical between WT and transgenic or gene-edited groups. We used the heteroscedastic standardized mean difference (SMD) as the effect size for the meta-analysis, employing random-effect models [80] (*Note 5 in Appendix2*). We estimated the overall mean effect sizes with 95% confidence intervals (CIs) and 95% prediction intervals (PIs) using the rma.mv function from the *metafor* package [81]. This allowed for the comparison of WT versus transgenic or gene-edited groups across parameters for each trait performance. Mean effect size estimates and their 95% CIs were reported, except where otherwise specified. Furthermore, SMD was interpreted using a rule of thumb similar to Cohen’s *d* but with some modifications: 0 < | SMD | ≤ 0.5, small effect; 0.5 < | SMD | ≤ 0.8, medium effect and | SMD | > 0.8, large effect [82].

### 2.5. Moderator analysis

We first created a null model to determine the overall effect of transgenesis or gene editing on animal trait enhancement. We included the effect size in the null model and random effects of: effect size ID, study ID and species name to control for non-independence of effect sizes. The *I*^2^ statistic (total heterogeneity) [83] was used to calculate the percent variance owing to inconsistencies in the population effect across studies, and *I*^2^ > 50% indicated significant between-study heterogeneity. We also quantified partial heterogeneity explained by each potential moderator using the function *i2*_*ml* from the *orchard* package [84]. The full model included moderators of taxon, parameter, method, species, gene, promoter, tissue, pathogen, generation, and phylogenetic relatedness as random effects. We also constructed a variance-covariance (VCV) matrix of species-phylogenetic relatedness to control for non-independent variation taken from the same study individuals for each dataset. We first conducted meta-regressions (full models) to determine how moderators affect the trait enhancement (growth, disease resistance and fatty acid content) led by transgenesis and gene editing. Then we established several meta-regressions to investigate the effect of each moderator (*Note 6 in Appendix2*). We calculated the total heterogeneity (*Q*_M_) and the proportion of total heterogeneity explained by moderators (marginal *R*^2^), using function *r2*_*ml* in the orchard package [84].

Specifically, we initially performed a method-moderator analysis to compare the effects of different methods (transgenesis *vs*. gene editing) on trait enhancement in animals, incorporating all parameters for each trait performance. Subsequent moderator analyses were conducted based on specific traits. For gene/protein expression, we assessed the overall effects of transgenesis and gene editing using a global model (all data) based on a method-moderator test. A taxon-moderator analysis was also conducted to assess expression profiles across taxa for both transgenesis and gene editing. Differences in gene/protein expression were examined across traits using trait-specific moderator analyses. In addition, gene- and tissue-moderator analysis were performed on both transgenesis and gene editing datasets to investigate if the expression of mRNA and protein was gene- and tissue-specific. Likewise, several moderators were analyzed in the growth traits, where a method-moderator analysis was used to compare the effects of transgenesis and gene editing on enhancing growth traits across different parameters. This was followed by taxon- and gene-moderator analysis to determine the effects across taxonomic classes and genes. The most promising candidate transgenes (coupled with promoters) and innate growth-regulating genes were identified through gene- and promoter-moderator analyses.

Regarding disease resistance and fatty acid traits, the positive and negative effect sizes jointly led to a null effect result when we combined all parameter datasets from transgenic and gene editing. To address this, we first performed parameter-moderator analysis to compare the effect of transgenesis and gene editing on individual traits across parameters. A taxon-moderator analysis was conducted for specific growth traits to assess the impact of transgenesis or gene editing on CSR, virus RNA, pathogen load and virus titer across taxonomic classes. Gene- and promoter-moderator analyses were also performed to confirm the valid candidate transgenes (coupled with promoters) and innate genes for disease resistance enhancement. Additionally, the effects of transgenesis and gene editing on improving disease resistance were evaluated through pathogen-moderator analysis. For the fatty acid traits, the available dataset of gene editing was limited to a single study by Park et al. [56], and all moderator-analysis were carried out using the transgenesis dataset, without conducting a method-moderator analysis. Taxon-moderator analyses were performed across parameters, such as ω-3, ω-6, the ratio of ω-6/ω-3, DHA and EPA to evaluate the effect of transgenesis on various fatty acid traits. The most effective candidate transgenes for fatty acid enhancement were proposed based on gene-moderator analyses. Lastly, generation-moderator analyses were performed for growth, disease resistance and fatty acid traits to investigate whether the improved traits caused by transgenesis or gene editing were similarly effective in offspring. Meta-regressions were generated using the *metafor* package to investigate how age influenced the effects of transgenesis and gene editing on growth and fatty acid traits, given the high degree of partial heterogeneity observed in these two traits.

### 2.6. Publication bias

To assess publication bias, funnel plots were generated, and all estimates were subjected to a classic Egger’s regression test (weighted regression with multiplicative dispersion) for funnel plot asymmetry [85]. A *P*-value < 0.05 implies that the funnel plot is asymmetric, and publication bias is possible. In this case, ‘trim and fill’ approach [86] was used to determine whether random-generated studies were needed to reduce potential publication bias. Finally, we tested for a time-lag bias for each trait performance to determine if effect sizes decline with more recent publication years [85].

### 2.7. Sensitivity analysis

We identified potential outliers and influential observations of the estimated measures across studies using sensitivity analyses to assess the stability and reliability of our meta-analysis. We performed a “leave-one-out” analysis for each trait, where one study was consecutively removed from the dataset and a new global meta-analytic mean and 95% was calculated [87].

## 3. Results

### 3.1. Data overview

The current meta-dataset extracted 171 figures from 154 studies (86 for transgenesis, 68 for gene editing) that provided relevant data on the effects of genetic engineering and gene editing on trait performance. In total, we obtained 1,145 effect sizes (SMD) across 10 taxonomic classes and 72 species/breeds, with the majority of studies conducted on livestock and ray-finned fishes (77 and 68 studies, respectively) (Fig. 1AB). A total of 55 genes were linked to the enhancement of target traits, of which 23 genes were associated with improved growth traits (15 for transgenesis, 8 for gene editing), 27 genes related to enhanced disease resistance (18 for transgenesis, 9 for gene editing), and 5 genes involved in altering fatty acid content (4 for transgenesis, 1 for gene editing) (Fig. 1C).

**Fig. 1.**
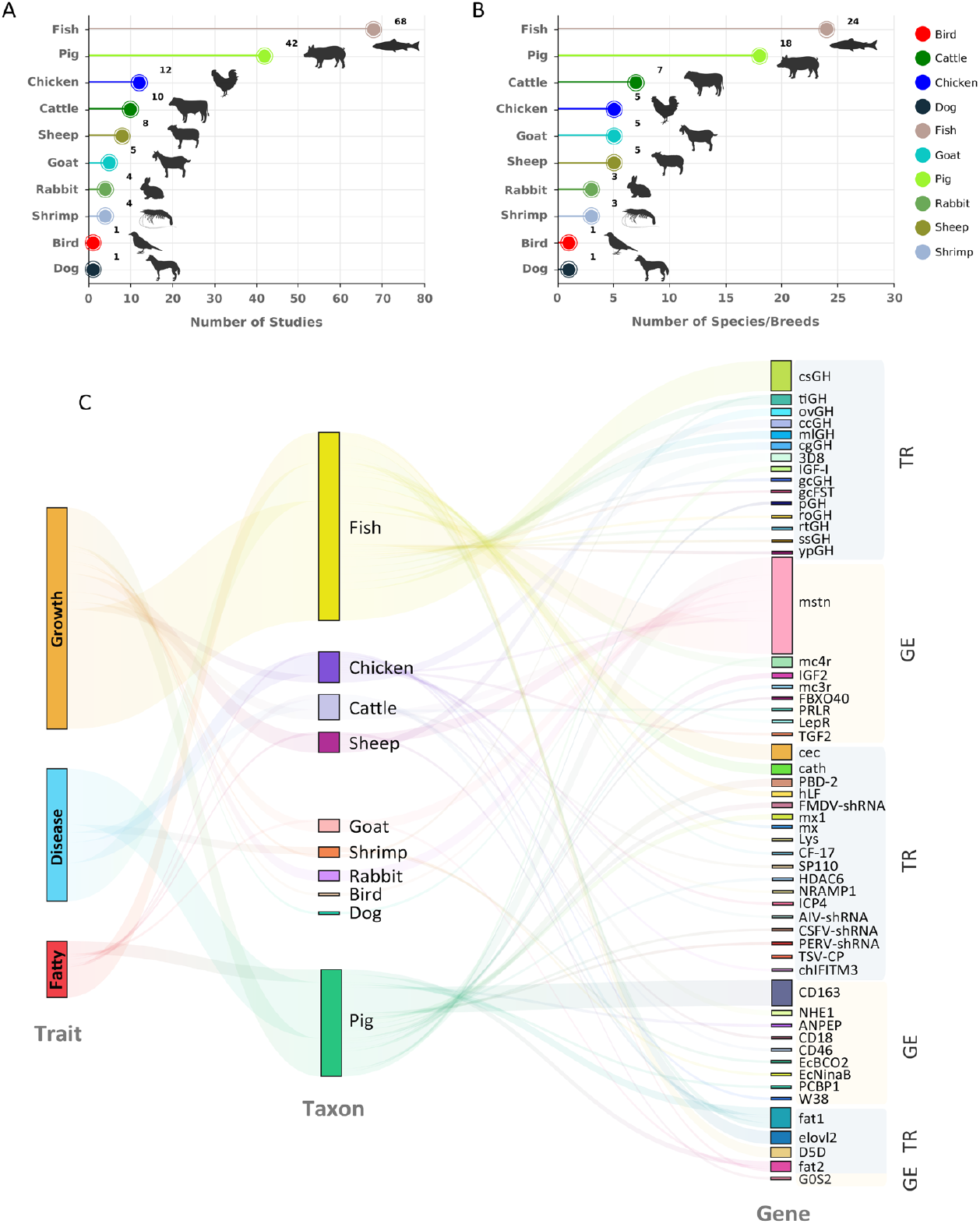
A summary of the current applications of transgenesis and gene editing for trait enhancement in farmed animals. **A** The number of studies focused on trait enhancement (growth, disease resistance and fatty acid) across taxa using transgenesis and gene editing. **B** The number of species/breeds involved in each taxonomic class. **C** The Sankey diagram shows the proportion of studies focused on different traits per taxon, and the target transgenes or innate genes are listed in the right column. TR, transgenesis; GE, gene editing.

### 3.2. Effects of transgenesis/gene editing on gene/protein expression

First, we assessed the expression profiles (both mRNA and protein) of target transgenes and innate genes at a global level by combing all data. Our multivariate meta-analysis model showed that there was moderate effect of transgenesis/gene editing on gene/protein expression (mean [95% CI]: 0.376 [−0.536 to 1.289], *t*_98_ = 0.819, *P* = 0.4149). The heterogeneity of our dataset was high (*I*^2^ = 100%), with 38% attributed to differences between studies, and 17.2% to differences between effect sizes. Notably, most effect sizes were derived from fish and pig species, and phylogenetic relatedness (*Fig. 2A in Appendix2*) explained 44.8% of heterogeneity, suggesting a phylogenetic effect on gene/protein expression.

**Fig. 2.**
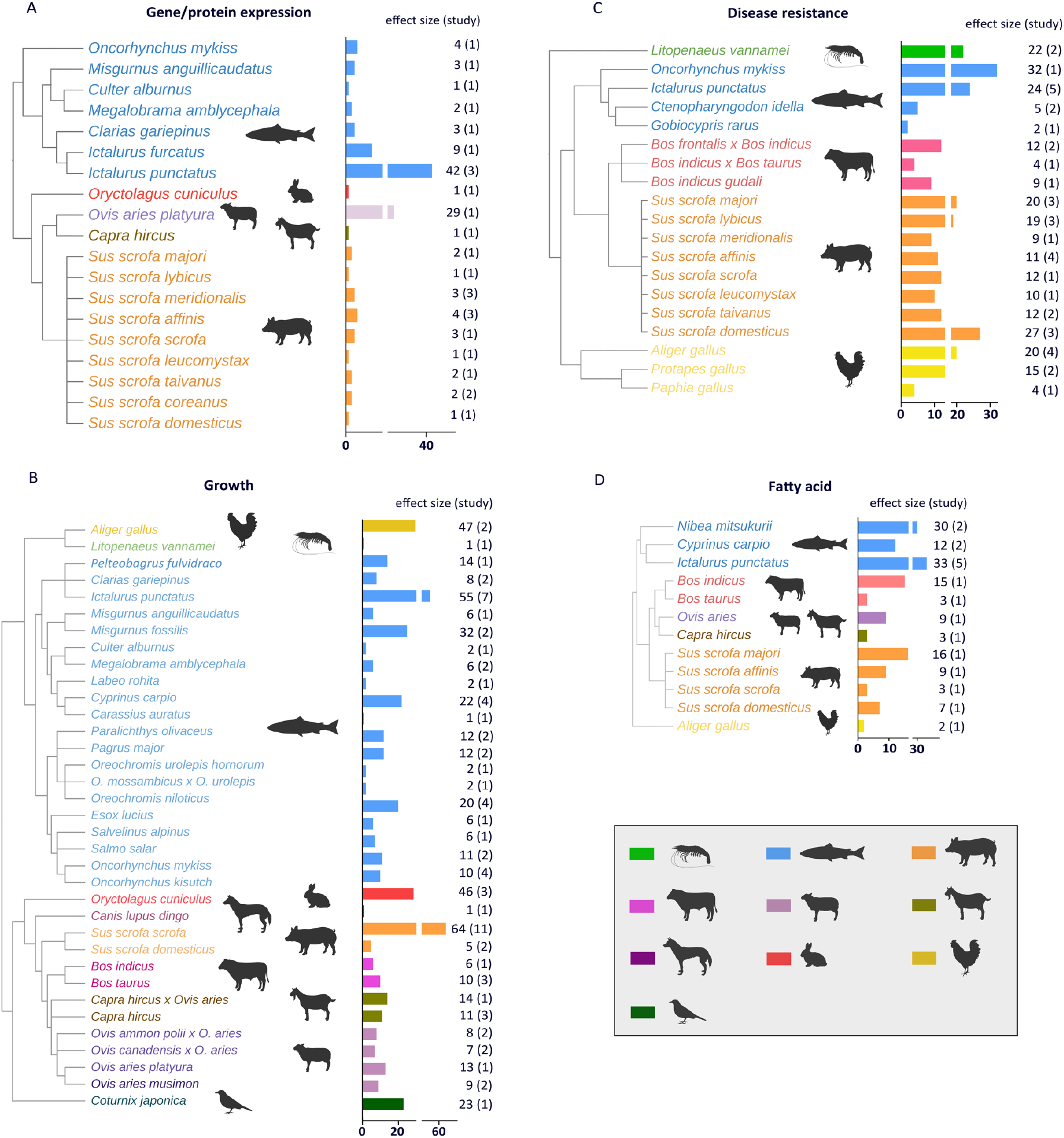
Phylogenetic tree of all species included in the gene expression (A), growth (B), disease resistance (C), and fatty acid (D) datasets. Colors represent taxonomic class. Bars and first column of numbers represent the number of effect sizes followed by the number of studies in brackets.

Obviously, transgenesis and gene editing significantly induced (mean [95% CI]: 4.876 [3.590 to 6.163], *P* < 0.0001) and reduced (mean [95% CI]: −3.990 [−5.789 to −2.192], *P* < 0.0001) mRNA/protein expression, respectively (*Fig. 3A in Appendix2*). Taxonomic class explained a large proportion of heterogeneity (*R*^2^ = 0.723, *Q*_M_ = 4.763, *P* = 0.9421, *df* = 11), when effects were averaged across all taxa. Specifically, transgenesis significantly increased the mRNA level and protein expression of the transgenes in different taxa (pig, mean [95% CI]: 5.653 [3.541 to 7.764], *P* < 0.0001 > sheep, mean [95% CI]: 4.953 [−0.542 to 10.447], *P* = 0.0773 > fish, mean [95% CI]: 3.336 [0.686 to 5.985], *P* = 0.0136) (*Fig. 3B, Fig. S2A in Appendix2*) in all three traits, including growth (mean [95% CI]: 4.701 [0.812 to 8.590], *P* = 0.0178), disease resistance (mean [95% CI]: 5.387 [2.951 to 7.823], *P* < 0.0001) and fatty acid (mean [95% CI]: 4.072 [1.007 to 7.137], *P* = 0.0092) (*Fig. 3C*). The expression of transgenes and proteins was gene- (*R*^2^ = 0.472, *Q*_M_ = 61.667, *P* < 0.0001, *df* = 10), promoter- (*R*^2^ = 0.265, *Q*_M_ = 22.140, *P* = 0.0011, *df* = 6) and tissue-specific (*R*^2^ = 0.263, *Q*_M_ = 25.210, *P* = 0.1537, *df* = 19) (*Fig. S2B-D in Appendix2*). Compared to other promoters, H1 exhibited the greatest induction of gene expression (mean [95% CI]: 12.618 [2.662 to 22.574], *P* = 0.0130), followed by β-actin and U6 (mean [95% CI]: 7.612 [4.258 to 10.967], *P* < 0.0001 for β-actin; mean [95% CI]: 4.562 [−4.114 to 13.239], *P* = 0.0863 for U6), while CMV and UBI promoters showed similar effects on gene expression (*Fig. S2E in Appendix2*). At the global level, these transgenes showed constant expression in both P1 and F1 generations (*Fig. S2F in Appendix2*).

**Fig. 3.**
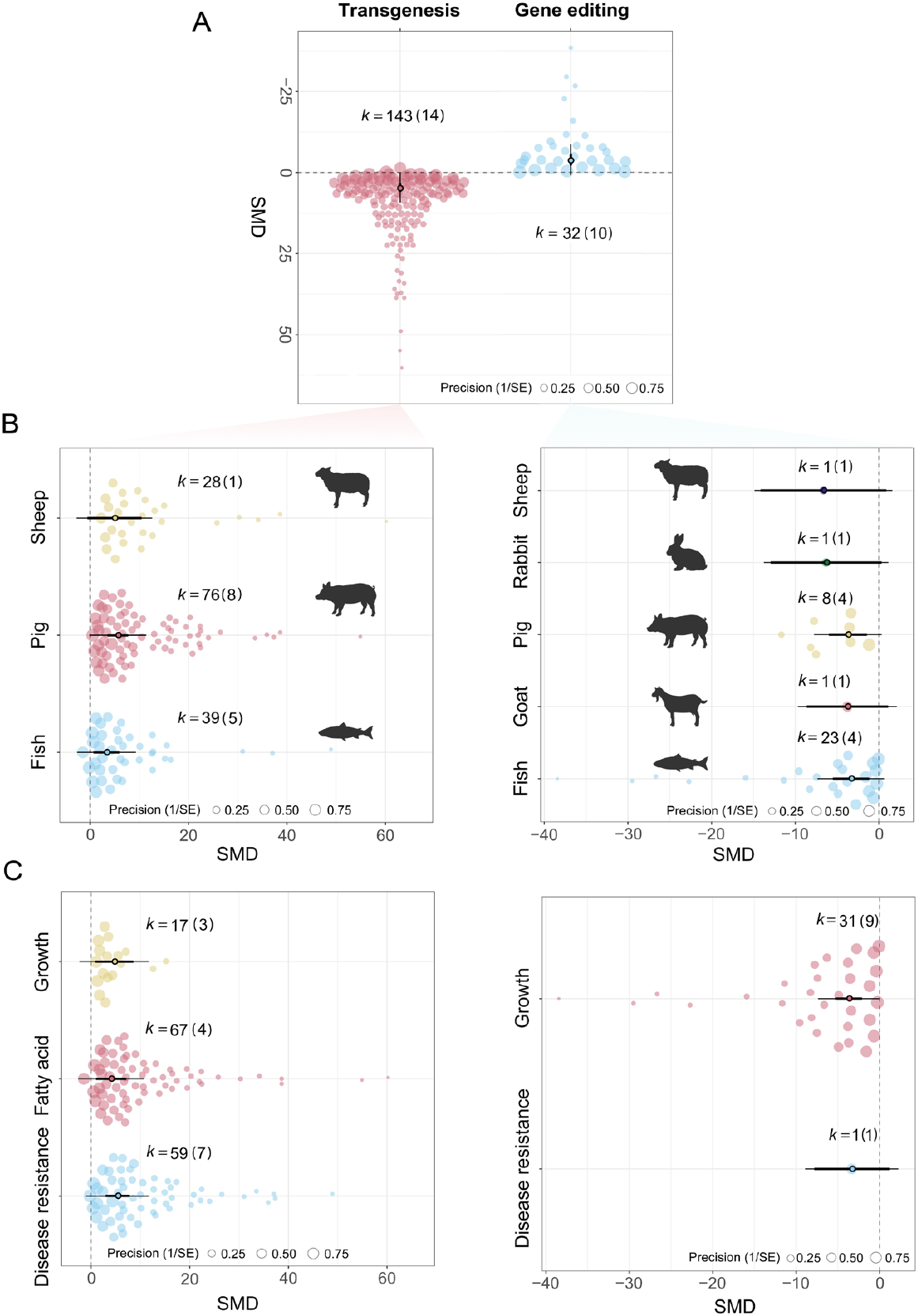
Significant induction of transgenes and inhibition of innate genes were observed across taxonomic classes and traits. **A** Overall effects of transgenesis and gene editing on gene/protein expression in the global model based on a method-moderator analysis. **B** Effect of transgenesis and gene editing on gene/protein expression for each taxon. **C** Effect of transgenesis and gene editing on gene/protein expression for each trait performance. The *x*-axis shows the values of effect sizes as heteroscedastic standardized mean difference (SMD), while the *y*-axis represents the density distribution of effect sizes. The size of the dots represents the precision of each effect size (1/SE). Thick black whisker lines and thin black lines represent 95% confidence intervals (CIs) and 95% prediction intervals, respectively. A bold error bar (95% CI) shows whether the overall effect size is significantly (*P* < 0.05) different from zero (i.e. not overlapping zero). *k* is the number of effect sizes and the number of studies (the number in brackets).

Gene editing had a comparable effect on gene/protein expression to transgenesis, as indicated by the nearly equal absolute values of the overall effect size (3.91 *vs*. 4.60) (*Fig. 3A*). Gene editing effectively disrupted gene/protein expression (*Fig. S3A in Appendix2*), and significant down-regulated expression of target innate genes/proteins was observed in fish and pig (mean [95% CI]: −2.834 [−5.238 to −0.430], *P* = 0.0209 for fish; mean [95% CI]: −3.692 [−5.922 to −1.462], *P* = 0.0012 for pig) (*Fig. 3B*). Similar to transgenesis, the moderator-analysis showed that the expression of edited innate genes was gene- (*R*^2^ = 0.007), species- (*R*^2^ = 0.519) and tissue-specific (*R*^2^ = 0.821) (*Fig. S3B-D in Appendix2*). Although 18 genes were edited across 42 animal species/breeds, expression data primarily came from traits related to growth and disease resistance (*Fig. 3C*), particularly for *mstn, IGF2, LepR* and *CD163* (*Fig. S3CE in Appendix2*), while expression data for other genes were seldom recorded.

### 3.3. Effects of transgenesis/gene editing on growth

Altered gene/protein expression resulting from transgenesis or gene editing led to enhanced trait performance, though these improvements were not consistent across traits and taxa, according to our meta-analysis. With respect to growth, we found significant effects of transgenesis/gene editing on growth in our full model, which included all moderators and parameters (mean [95% CI]: 0.337 [0.230 to 0.444], *t*_499_ = 6.213, *P* < 0.0001). Heterogeneity in this dataset was high (*I*^2^ = 99.8%), with 16.8% attributed to true differences between studies, 48.2% to differences between effect sizes, and phylogenetic relatedness (*Fig. 2B*) explained 34.9% of heterogeneity.

Both transgenesis and gene editing had significant positive effects on growth enhancement, either by integrating foreign genes or disrupting innate growth-regulating genes (mean [95% CI]: 1.626 [0.679 to 2.572], *k* = 172, *P* = 0.0008 for transgenesis; mean [95% CI]: 2.009 [1.141 to 2.877], *k* = 343, *P* < 0.0001 for gene editing) (*Fig. S4A in Appendix2*). This improvement in growth traits was confirmed by changes in several parameters, including increased body weight, weight gain, SGR, plasma GH and decreased FCE (Fig. 4A). Although large significant effects were detected in the P1 generation, the F1 generation inherits these effects with smaller effect sizes compared to its parents (mean [95% CI]: 4.465 [2.706 to 6.225], *k* = 48, *P* < 0.0001 for P1; mean [95% CI]: 1.205 [−0.507 to 2.918], *k* = 124, *P* = 0.1677 for F1) (*Fig. S4B in Appendix2*). Taxon explained a small proportion of heterogeneity (*R*^2^ = 0.021, *Q*_M_ = 1.528, *P* = 0.6757, *df* = 3), and significant positive effects of transgenesis on growth were primarily observed in fish (mean [95% CI]: 1.982 [0.207 to 3.756], *k* = 172, *P* = 0.0287). Although small and large effects were noted in transgenic pig and sheep, respectively, these differences were not statistically significant compared to the WT group (mean [95% CI]: −0.254 [−6.099 to 5.591], *k* = 9, *P* = 0.9320 for pig; mean [95% CI]: 3.998 [−1.947 to 9.942], *k* = 23, *P* = 0.1875 for sheep) (*Fig. S4C in Appendix2*). In addition, 9 (*mlGH, cgGH, ovGH, roGH, csGH, ssGH, tiGH, gcFST* and *rtGH*) out of 15 GH transgenes showed large positive effects on growth enhancement, with *mlGH, cgGH*, and *ovGH* being the most impactful (*Fig. S4D in Appendix2*). Compared to transgenesis, gene editing exhibited a larger effect on growth enhancement as reflected by a larger effect size (2.01 *vs*. 1.63) (*Fig. S4A in Appendix2*). Gene editing significantly improved growth traits with large effects in 5 of 10 taxonomic classes, including cattle, fish, goat, pig and sheep (mean [95% CI]: 2.007 [0.386 to 3.628], *k* = 14, *P* = 0.0152 for cattle; mean [95% CI]: 1.898 [1.221 to 2.575], *k* = 113, *P* < 0.0001 for fish; mean [95% CI]: 2.073 [1.204 to 2.943], *k* = 25, *P* < 0.0001 for goat; mean [95% CI]: 1.865 [1.054 to 2.676], *k* = 56, *P* < 0.0001 for pig; mean [95% CI]: 3.023 [1.525 to 4.521], *k* = 15, *P* < 0.0001 for sheep). Inexplicably, gene editing showed a weak effect on chicken growth (mean [95% CI]: 0.051 [−1.932 to 2.033], *P* = 0.9601) even when a large sample size (*k* = 46 > 30) of effect sizes was collected (*Fig. S4E in Appendix2*). Gene editing showed broader impacts on growth traits compared to transgenesis, significantly improving a wider range of growth parameters, such as condition factor (mean [95% CI]: 3.098 [2.527 to 3.670], *k* = 13, *P* < 0.0001), myofiber number (mean [95% CI]: 2.493 [1.869 to 3.118], *k* = 27, *P* < 0.0001), and myofiber size (mean [95% CI]: 1.247 [0.685 to 1.808], *k* = 41, *P* < 0.0001), in addition to body weight, weight gain, and SGR (Fig. 2B). However, the meta-analysis indicated that gene editing had only a moderate, non-significant effect on FCE (mean [95% CI]: 0.759 [−0.042 to 1.561], *k* = 6, *P* =0.0634) (Fig. 4B). Although a total of 8 growth-regulating genes were targeted by gene editing tools to increase growth in farmed animals, the effects were variable across studies. Encouragingly, disruption of *mstn, mc4r* or *IGF2* showed a large significant positive effect on growth, followed by *PRLP, LepR* and *mc3r*. There were no significant general effects on animal growth when the *TGFβ2* or *FBXO40* was edited (*Fig. S4F in Appendix2*).

**Fig. 4.**
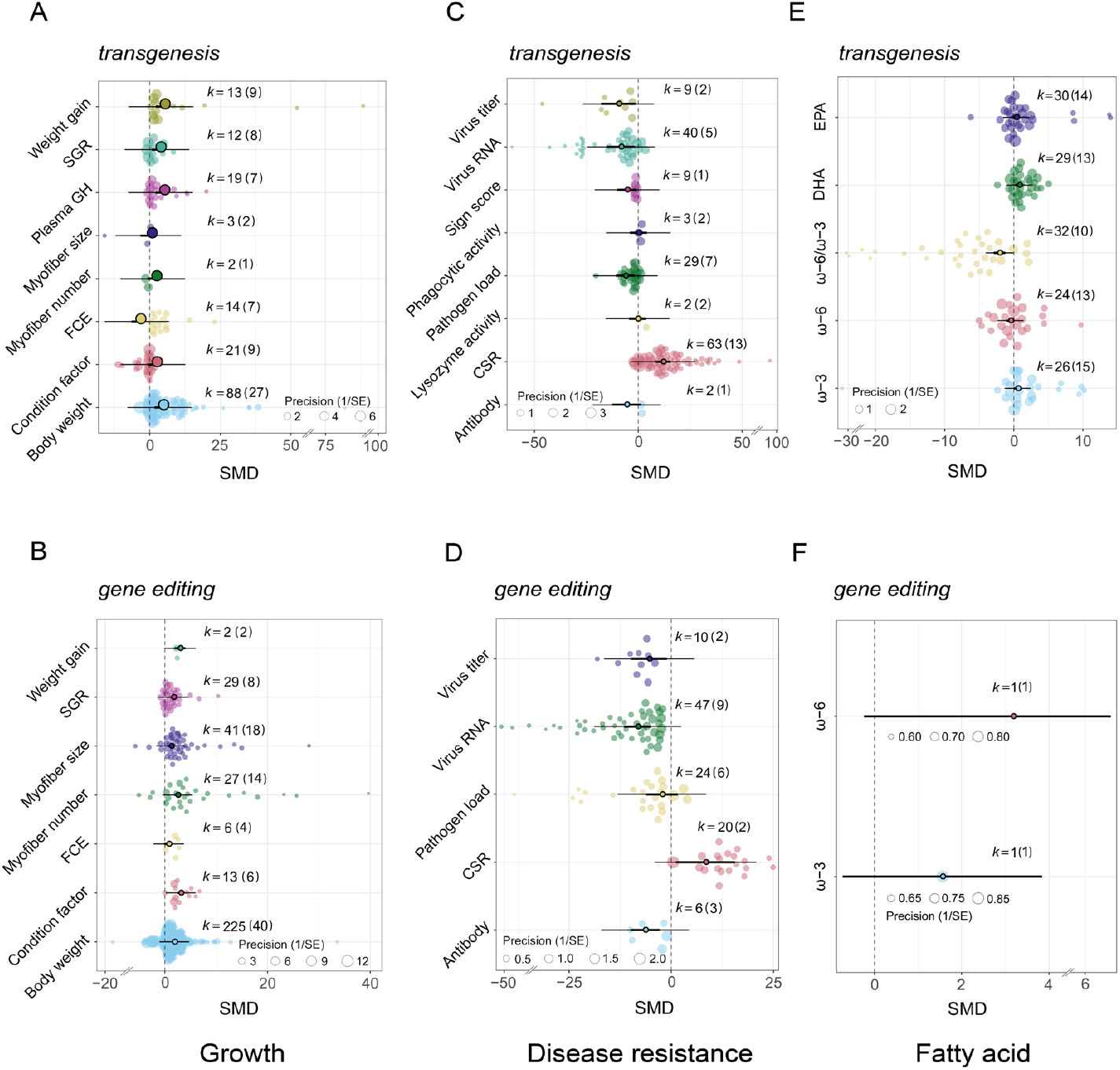
Comparison of effects of transgenesis and gene editing on trait enhancement in farmed animals across various parameters. **A** Effect of transgenesis on individual growth traits across 19 species/breeds in the meta-dataset. **B** Effect of gene editing on individual growth traits in 38 species/breeds. **C** Effect of transgenesis on disease resistance traits in 16 species/breeds. **D** Effect of gene editing on disease resistance traits across 7 species/breeds. **E** Effect of transgenesis on fatty acid traits across 10 species/breeds. **F** Effect of gene editing on fatty acid traits in White Leghorn chicken.

We found evidence for an age-dependent effect of transgenesis and gene editing on growth traits. The effect of transgenes or edited genes on growth enhancement decreased with age. This result was mainly supported in SGR and myofiber (number and size), but not in body weight (Fig. 5A).

**Fig. 5.**
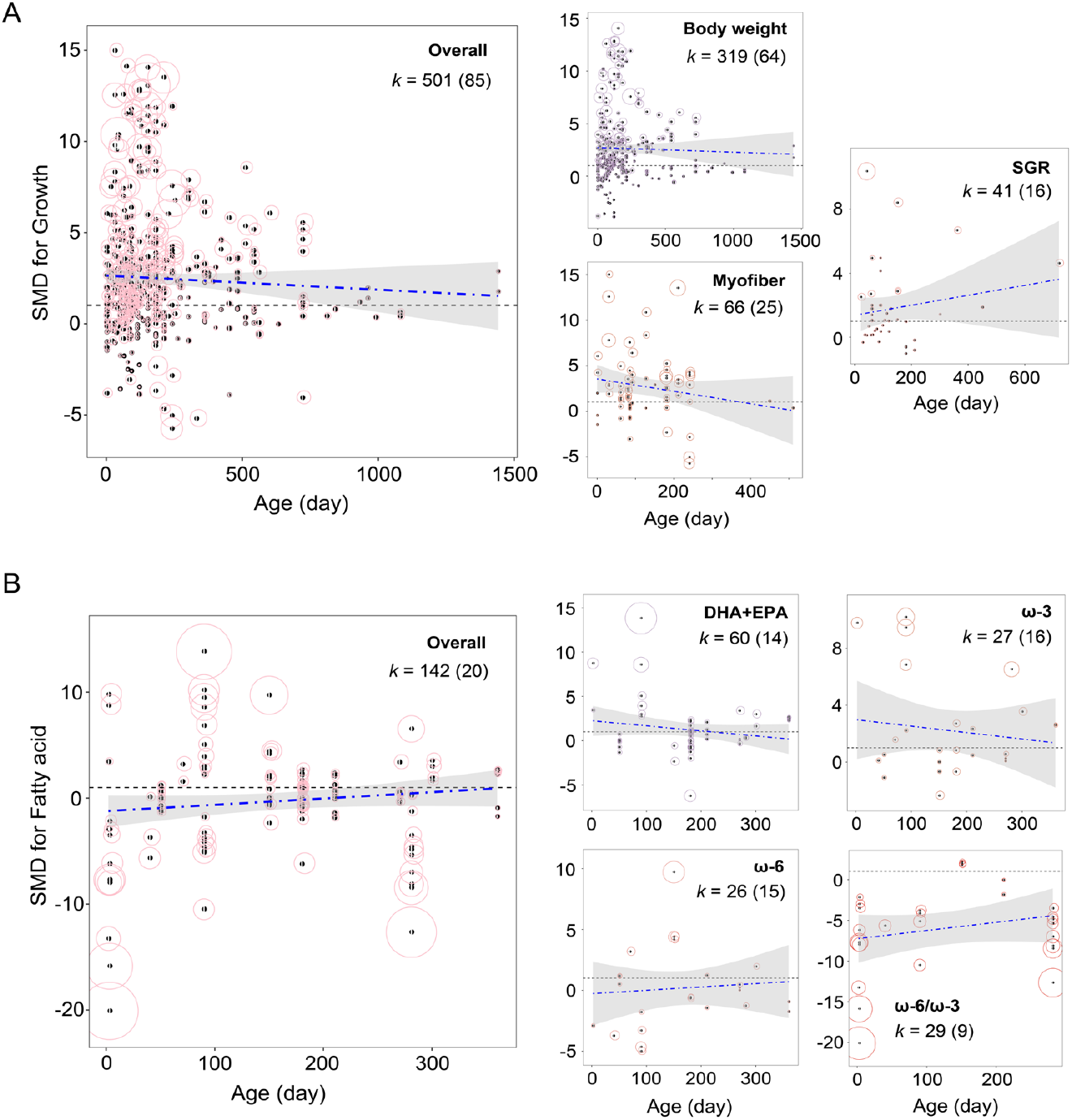
The impact of transgene integration or gene knockout on traits progressively decreases with advancing age. **A** Effects of age (*x*-axis) on the effect size (SMD) (*y*-axis) of growth traits across the entire dataset, and disaggregated for body weight, myofiber and SGR. **B** Effects of age (*x*-axis) on the effect size (SMD) (*y*-axis) of fatty acid traits across the entire dataset, and broken down for DHA+EPA, ω-6, ω-3 and ω-6/ω-3. The center of each circle indicates the mean effect size. The diameter of circles represents 95% confidence interval (CI) of the effect size. The dark line with shaded bars shows the overall effect of age on effect sizes and its 95% CI, respectively, and the blue dotted line shows the mean regression line. *k* is the number of effect sizes and the number of studies (the number in brackets).

### 3.4. Effects of transgenesis/gene editing on disease resistance

We did not find a significant general effect of transgenesis/gene editing on growth using a full model (including all moderators and parameters; mean [95% CI]: −0.373 [−0.887 to 0.140], *t*_235_ = −1.433, *P* = 0.1531). Heterogeneity in this dataset was still high (*I*^2^ = 100%), with 41.3% attributed to true differences between studies, 15.8% to differences between effect sizes, and phylogenetic relatedness (*Fig. 2C*) explained 42.8% of heterogeneity.

A total of 8 parameters (*Appendix1*) related to disease resistance traits were collected in our dataset. Parameter explained a large significant proportion of heterogeneity (*R*^2^ = 0.564, *Q*_M_ = 52.164, *P* < 0.0001, *df* = 7). The parameter-moderator analysis showed that transgenesis had various effects (both positive and negative) on these parameters, which caused a non-significant pooled effect size (*k* = 157, *P* = 0.0981). In this case, we first determined different effects of transgenesis on various parameters, including significant positive effects on CSR (mean [95% CI]: 11.928 [8.216 to 15.640], *k* = 63, *P* < 0.0001), and negative effects on pathogen load (mean [95% CI]: − 6.191 [− 10.671 to – 1.710], *k* = 29, *P* = 0.0068), sign score (mean [95% CI]: − 5.425 [− 10.051 to – 0.780], *k* = 9, *P* = 0.0215), virus RNA (mean [95% CI]: − 8.398 [− 15.212 to – 1.584], *k* = 40, *P* = 0.0157), and virus titer (mean [95% CI]: − 9.483 [− 17.806 to – 1.160], *k* = 9, *P* = 0.0255). However, transgenesis is less likely to affect antibody response (mean [95% CI]: − 5.650 [−12.568 to 1.268], *k* = 2, *P* = 0.1094), lysozyme activity (mean [95% CI]: − 0.299 [− 4.380 to 4.152], *k* = 2, *P* = 0.8859) and phagocytic activity (mean [95% CI]: 0.0641 [− 4.024 to 4.152], *k* = 3, *P* = 0.0001) (Fig. 4C).

The improvement in disease resistance through transgenesis varied across taxonomic classes depending on the specific parameters assessed. Specifically, a significant increase of CSR in fish (*Fig. S5A in Appendix2*), a substantial reduction of virus RNA levels in pig (*Fig. S5B in Appendix2*), and a significant decrease of pathogen load in both chicken and cattle (*Fig. S5C in Appendix2*) were observed. For the virus titer, a total of 9 effect sizes from chicken and pig were reported in our dataset, showing no general reduction in virus titer for either species (*Fig. S5D in Appendix2*). Our findings suggest that different transgenes, in combination with various promoters, exhibited diverse impacts on the enhancement of disease resistance traits. The top five genes (*CF-17, mx, cath, cec* and *NRAMP1*) (*Fig. S5EF in Appendix2*) driven by appropriate promoters (UBI, β-actin, CMV or U6) (*Fig. S5GH in Appendix2*) had the greatest effect on improving disease resistance.

Gene editing appeared to have a greater impact on disease resistance traits at a global level compared to transgenesis, as evidenced by a larger absolute effect size (mean [95% CI]: 1.268 [−2.061 to 4.597], *k* = 157, *P* = 0.4554 for transgenesis; mean [95% CI]: −3.912 [−8.080 to 0.257], *k* = 107, *P* = 0.0659 for gene editing) (*Fig. S6A in Appendix2*). Gene editing not only significantly enhanced CSR post-pathogen infection (mean [95% CI]: 8.355 [1.154 to 15.556], *k* = 20, *P* = 0.0230), and reduced virus RNA or titer levels (mean [95% CI]: − 8.234 [−11.557 to −4.911], *k* = 47, *P* < 0.0001 for virus RNA; mean [95% CI]: −5.414 [−9.790 to −1.039], *k* = 10, *P* = 0.0153 for virus titer), but also significantly inhibited antibody responses (mean [95% CI]: − 6.400 [−9.923 to −2.878], *k* = 6, *P* = 0.0004), a result not presented in the transgenesis dataset (Fig. 4D). Furthermore, the effects of gene editing on disease resistance enhancement varied across taxa (*R*^2^ = 0.678, *Q*_M_ = 38.560, *P* < 0.0001, *df* = 3). For instance, a significant reduction in virus RNA levels was determined in pigs but not in chicken or cattle (*Fig. S6B in Appendix2*), with *PCBP1* and *CD163* genes contributing to these effects (*Fig. S6C in Appendix2*). Notably, both transgenesis and gene editing showed variability in their effects across different pathogens (*Fig. S6DE in Appendix2*), with both approaches having the greatest inhibitory effects on bacteria, followed by viruses and parasites.

### 3.5. Effects of transgenesis/gene editing on fatty acid

The effects of transgenesis or gene editing on fatty acid composition showed no significant overall impact when using a full model that accounted for all moderators and parameters (mean [95% CI]: −0.09 [−0.392 to 0.218], *t*_140_ = −0.565, *P* = 0.5732). A high level of heterogeneity (*I*^2^ = 99.8%) was detected, with 12.6% attributed to true differences between studies, 87.2% to differences between genes, while we did not find phylogenetic relatedness (*Fig. 2D*), indicating the phylogenetic structure does not contribute to the observed variation in effect sizes.

A large proportion of heterogeneity in taxon and gene were observed, respectively, when effects were averaged across all parameters (*R*^2^_taxon_ = 0.591, *R*^2^_gene_ = 0.575). We performed taxon- and gene-mediated moderator-analyses for each parameter. Although published work reported significant improvements in ω-3, DHA, EPA and reductions in ω-6 and the ω-6/ω-3 ratio, our meta-analytic findings showed significant effects on ω-3, DHA, and ω-6/ω-3 (mean [95% CI]: 0.584 [−0.009 to 1.177], *k* = 26, *P* = 0.0537 for ω-3; mean [95% CI]: 0.792 [0.201 to 1.383], *k* = 29, *P* = 0.0086 for DHA; mean [95% CI]: −2.073 [−2.909 to −1.237], *k* = 32, *P* < 0.0001 for ω-6/ω-3), but no significant effect on EPA or ω-6 (mean [95% CI]: 0.342 [−0.246 to 0.930], *k* = 30, *P* = 0.2545 for EPA; mean [95% CI]: −0.471 [−1.059 to 0.117], *k* = 24, *P* = 0.1165 for ω-6) when all data were combined globally (Fig. 4E). These effects were taxon-specific across different parameters. The largest positive effect of transgenesis on ω-3 was observed in cattle, followed by pig and fish (mean [95% CI]: 4.671 [0.767 to 8.576], *k* = 4, *P* = 0.0190 for cattle; mean [95% CI]: 4.065 [0.657 to 7.474], *k* = 6, *P* = 0.0194 for pig; mean [95% CI]: 1.186 [−0.611 to 2.984], *k* = 14, *P* = 0.1959 for fish) (*Fig. S7A in Appendix2*). Similarly, the largest effect on ω-6, ω-6/ω-3, DHA and EPA were found in cattle, cattle, pig and pig, respectively. Nevertheless, we found no significant effect of transgenesis on the level of ω-3 in goat (mean = 0.099, *k* = 1, *P* = 0.9696), ω-6/ω-3 in fish (mean = 0.797, *k* = 2, *P* = 0.8845) or DHA in cattle (mean = 1.571, *k* = 3, *P* = 0.2560) (*Fig. S7B-E in Appendix2*). Different transgenes appeared to play distinct roles in enhancing fatty acid traits. Integration of *fat1* transgene significantly improved levels of ω-3, DHA and EPA (*Fig. S8A-C in Appendix2*), whereas decreasing ω-6 and ω-6/ω-3 (*Fig. S8D-E in Appendix2*). In contrast, transfer of *fat2* did not significantly increase ω-3 level or reduce ω-6, but it dramatically lowered the ratio of ω-6/ω-3. However, the *elovl2* (*k* = 6 for ω-3, *k* = 5 for ω-6, *k* = 15 for DHA, *k* = 15 for EPA, all *P* > 0.05) and *D5D* transgenes (*k* = 6 for ω-3, *k* = 6 for ω-6, *k* = 6 for DHA, *k* = 6 for EPA, all *P* > 0.05) were less likely to alter fatty acid traits across all taxonomic classes (*Fig. S8 in Appendix2*).

To date, only one gene editing-based study by Park et al. [56] involving one gene (*G0S2*) with two effect sizes has been reported in chickens for the enhancement of fatty acid. Although the study reported significant improvements in ω-3 and ω-6 levels, we found no significant effects on fatty acid alteration in *G0S2*-KO chicken compared to WT individuals (*k* = 1, *P* = 0.1813 for ω-3; *k* = 1, *P* = 0.1681 for ω-6) (Fig. 4F). Additionally, we found evidence that the effects of transgenesis and gene editing on fatty acid diminished with advancing age, as reflected by decreased effects on DHA+EPA, ω-3, ω-6, and ω-3/ω-6 (Fig. 5B).

### 3.6. Publication bias

We found little evidence of publication bias for gene/protein expression (*t*_173_ = 0.582, *P* = 0.5613) and fatty acid traits (*t*_133_ = 0.243, *P* = 0.8083) based on our results. The funnel plots and results of Egger’s regression test indicated potential publication bias for the effect sizes of growth (*t*_470_ = 5.274, *P* < 0.0001) and disease resistance (*t*_264_ = 5.179, *P* < 0.0001). In these cases, the trim-and-fill method was used to estimate the number of missing effect sizes needed for the current meta-analysis. The results revealed that 141 and 73 additional effect sizes, respectively, were needed to compensate for the potential publication bias (*Fig. S10 in Appendix2*). Nevertheless, transgenesis and gene editing still had significant effects on growth and disease resistance traits (adjusted mean [95% CI]: −1.249 [−1.516 to −0.982], *k* = 613, *P* < 0.0001 for growth; 2.953 [2.142 to 3.764], *k* = 339, *P* < 0.0001 for disease resistance) after incorporating these randomly generated effect sizes. We also did not detect a decline in the magnitudes of effects in more recent years (*Fig. S11 in Appendix2*).

### 3.7. Sensitivity analysis

The influence analysis indicated that no studies carried a significant deviation of pooled effect size from the overall level when we removed studies one by one (*Fig. S12 in Appendix2*). By examining the leave-one-out results, we confirmed that no single study dominated the results. Notably, the results from the full model were consistent regardless of whether the VCV matrix was included (gene/protein expression, disease resistance, and fatty acid), so we present the model without the VCV matrix throughout the main text.

## 4. Discussion

Advances in genetic engineering and genome editing technologies have led to consistent enhancement of trait performance in modified animals across individual studies. Here, we conduct a comprehensive meta-analysis quantifying the effects of transgenesis and gene editing on trait performance across various animal taxa. Our findings challenge the assumption that genetic modifications uniformly result in trait augmentation, showing considerable variability in outcomes such as growth, disease resistance, and fatty acid content. This variability is shaped by factors such as the specific animal taxa, the traits being targeted, and the transgenes or innate genes involved. In aggregate, this study reveals the effects of genetic modifications on trait enhancement across animals and highlights key gaps in knowledge that will facilitate a better understanding of the future applications of transgenesis and gene editing.

Our results indicate that both transgenesis and gene editing significantly impact key phenotypic traits, albeit with varying degrees of influence. Genetic modifications notably enhanced gene/protein expression, growth, and disease resistance, while their effects on fatty acid composition were more inconsistent. Specifically, gene expression was strongly affected by both methods, highlighting the ability of transgenesis or gene editing to effectively upregulate or downregulate target genes. Transgenesis generally resulted in higher gene expression by introducing new functional genes under robust promoters like H1 or *β*-actin, whereas gene editing often downregulated or knocked out specific genes, particularly growth regulators such as *mstn* and *IGF2*, illustrating the precision of gene pathway manipulation. This modulation of gene expression led to marked improvements in growth traits, including body weight and SGR, with gene editing showing a greater effect size for growth compared to transgenesis, likely due to its precise targeting of negative growth regulators with low off-target events [1]. These enhancements were especially pronounced in fishes, likely reflecting differences in genetic architecture and growth physiology across taxa [57,58]. In terms of disease resistance, gene editing demonstrated more consistent and robust effects than transgenesis, as evidenced by larger effect sizes and broader impacts across trait parameters. Our analysis revealed that targeted gene knockdown effectively enhanced disease resistance in livestock by significantly reducing the incidence of infectious diseases through the suppression of viral receptors and immune regulatory genes. However, the variability across species and pathogens indicates that further optimization is needed, particularly in transgenesis, where the overall effects on disease resistance were not significant. Additionally, fatty acid composition showed less consistent changes, with no significant overall effect detected; however, improvements in specific parameters, such as the ω-6/ω-3 ratio and DHA levels, were observed in certain species like pigs and cattle. The variability in fatty acid traits suggests that while genetic manipulation of lipid metabolism is possible, its success may depend on factors such as species, specific transgenes or edits, and their integration into existing metabolic pathways [11,52,53]. These findings highlight the advantages of gene editing over transgenesis due to its precision, efficiency, and ability to produce natural-like modifications, suggesting greater acceptability in regulatory and consumer contexts, while also serving as valuable references for government regulation of transgenic and gene-edited animals.

This meta-analysis highlights significant taxon-specific variation across parameters in responses to genetic interventions, with fish showing the strongest improvements, particularly in growth. This may be due to the focus on introducing a diversity of *GH* genes in fishes [23,59], which are known to enhance growth rate and body size. In contrast, mammalian species like pigs, sheep, and cattle displayed more variable effects, reflecting differences in genetic regulation of traits [7,60] like growth and disease resistance. The variability in effect sizes across traits underscores the importance of targeting specific genes or pathways. Growth traits consistently improved across taxa due to simpler interventions, while disease resistance showed more variability, likely due to the complexity of immune pathways [61,62]. The success of genetic interventions was also influenced by promoter type, with the H1 promoter driving the most effective gene expression, followed by β-actin and U6, emphasizing the need for careful promoter selection to optimize transgenesis.

In our meta-dataset comprising animals of various ages, we observed that the effects of transgenesis and gene editing on growth and fatty acid traits varied with age. Specifically, there was an age-dependent reduction in the efficacy of these modifications, suggesting their impact diminishes over the animal’s lifespan. This may result from diminished cellular activity, metabolic slowdown, hormonal changes, and compensatory mechanisms [59,63]. Furthermore, trait improvements were more pronounced in the P1 generation than in F1. Despite consistent expression or repression of transgenes or edited genes in F1 generation (*Fig. S2F, Fig. S9A in Appendix2*), the effects on growth (*Fig. 5B, Fig. S9B in Appendix2*), disease resistance (*Fig. S9CD in Appendix2*), and fatty acids (*Fig. S9E in Appendix2*) were less marked in offspring. This could be due to partial dominance or incomplete penetrance, where full expression requires homozygosity, yet the F1 generation, often heterozygous [43,46,49,64,65], exhibits reduced trait manifestation. Epigenetic modifications or other intergenerational regulatory mechanisms may also play a role [66]. To more fully assess these effects, future analyses should include homozygous animals beyond the F1 generation.

Our meta-analysis identified 18 innate genes involved in trait enhancement, with only 5 showing significant global-level improvements. For instance, disruption of *mstn, mc4r*, or *IGF2* genes significantly enhanced growth in both livestock and fish, while knockouts of *CD163* or *PCBP1* increased viral resistance by reducing viral RNA in livestock. Despite this, few genes have been validated for trait improvement in farm animals, particularly in fish, posing a significant challenge for genome manipulation aimed at enhancing economically important traits. The limited number of identified genes linked to such traits constrains the application of genetic manipulation techniques, including gene editing [67,68]. Therefore, focused efforts on identifying specific gene variants through comprehensive genomic studies— such as GWAS, QTL mapping, and functional genomics—are critical [68]. Once identified, targeted genetic manipulations can be applied to introduce beneficial variants or remove deleterious ones, improving the predictability and effectiveness of breeding programs.

As with any study, limitations must be acknowledged, and research gaps remain. In this study, we analyzed and interpreted effect sizes across taxa, traits, parameters, and gene-specific factors, employing species-moderator analysis rather than phylogenetic meta-analysis due to low variation across species. Our literature screening may have overlooked studies, as our meta-dataset only includes publications in English. Additionally, we excluded studies on model animals (i.e., zebrafish, medaka and mice) [69] and *in-vitro* studies identifying potential causative genes using cell lines. For example, several representative genes from fishes, like rhamnose-binding lectin (*RBL*), signal transducer and activator of transcription 2 (*STAT2*), junctional adhesion molecule-A (*JAM*-*A*), the repressor of RNA polymerase III transcription of *Paralichthys olivaceus* (*PoMaf1*), NEDD-8 activating enzyme 1 (*nae1*), and GRB2-associated binding protein 3 (*GAB3*) have undergone induced mutations that mimicked and altered the immunity of the fish and improved the host’s resistance to disease [70,71]. Therefore, our results represent broad trends from the current literature that meet our inclusion criteria and should be interpreted accordingly.

Future primary studies should aim to cover a broader range of taxonomic groups (i.e., model animals in the laboratory) and traits (i.e., reproductive outcomes) that were underrepresented in our dataset. It would be interesting to examine the effects of genetic modification on model animals beyond livestock and fish, which were not included in this study, to uncover the process of transgenesis and gene editing from model to captive animals. Reproduction confinement is another aim of transgenesis and gene editing in farm animals [7,68], and a growing number of studies have identified candidate reproduction-regulating genes, particularly in fish species [68]. It would also be valuable to verify these promising genes by assessing reproductive outcomes (i.e., gamete quality, sperm motility, fertilization rate, hatching rate) on a global scale for transgenic or gene-edited animals.

## Acknowledgements

L.L. was supported by a Ten Thousand Talent Plans for Young Top-notch Talents Grant (20221116). D.R. was supported by a Horizon Europe Framework Programme Grant (101079467).

## Author contributions

J.W., S.N., and L.L.: Designed the study. J.W., J.W., L.L., R.S., A.F.: Collected the data. S.N., R.A.D. and D.R.: Provided extra studies. L.L.: Checked for repeatability of data extraction. J.W., S.N., F.L., G.W and L.L.: Performed the meta-analyses. J.W., L.L.: Wrote the first draft of the manuscript, and all authors contributed substantially to revisions.

## Competing interests

The authors declare no competing interests.

## Notes

### Competing Interest Statement

The authors have declared no competing interest.

